# Translocation of structured biomolecule through a vibrating nanopore

**DOI:** 10.1101/296780

**Authors:** M. A. Shahzad

## Abstract

We study the effect of fluctuating environment in protein transport dynamics. In particular, we investigate the translocation of a structured biomolecule (protein) across a temporally modulated nano-pore. We allow the radius of the cylindrical pore to oscillate harmonically with certain frequency and amplitude about an average radius. The protein is imported inside the pore whose dynamics is influences by the fluctuating nature of the pore. We investigate the dynamic and thermodynamical properties of the translocation process by revealing the statistics of translocation time as a function of the pulling inward force acting along the axis of the pore, and the frequency of the time dependent radius of the channel. We also examine the distribution of translocation time in the intermediate frequency regime. We observe that the shaking mechanism of pore leads to accelerate the translocation process as compared to the static channel that has a radius equal to the mean radius of oscillating pore. Moreover, the translocation time shows a global maximum as a function of frequency of the oscillating radius, hence revealing a resonant activation phenomenon in the dynamics of protein translocation.

## I. INTRODUCTION

Translocation of biomolecule play an important role to understand the DNA sequencing and sorting, and drug delivery mechanism. Translocation of DNA through biological pore was first performed using voltage-driven translocation experiment. In this device, a constant voltage bias, usually of the order of magnitude of mV, generate an ionic current through the channel that can be recorded by standard electro-physiological technique like patch-clamp. One bio-polymer is introduced on the cis-side, it crosses the pore driven by thermal fluctuation and by the voltage and flows into the opposite chamber, the trans-side. Its passage temporarily clogs the channel and provokes a detectable ion current drop, which strongly depends on the chemical and physical properties of the molecule that occupied the pore. Moreover, the duration of current loop is a direct measure of the translocation time, which is in turn used to characterize a single transport event, along with the intensity of the blockage itself [1–9]. Nanopores have become an important tool for molecule detection at single molecular level. With the development of fabrication technology, synthesized solid-state membranes are promising candidate substrates in respect of their exceptional robustness and controllable size and shape. It opens a new possible way to read out the sequence of DNA without amplification and labeling [10, 11]. Due to its controllable pore size, and shape, long term stability and ease of integration into detection devices, solid-state nano-pores can serve as a multifunctional sensor for detection and characterization of protein [12, 13], DNA/protein [14] or nano-materials/nano-particles [15, 16].

ATP-dependent proteases of the AAA+ (ATPases associated with various cellular activities) power the degradation of abnormal, denatured, or otherwise damaged polypeptide, as well as the removal of short-live regulatory proteins [23]. ClpX is a component of the ClpXP proteasome-like complex that is responsible for the targeted degradation of numerous protein substrate in Escherichia coli and other organisms. Within this complex, ClpX forms a homohexameric ring that uses ATP hydrolysis to unfold and translocate proteins through its central pore and into a proteolytic chamber (ClpP) for degradation.

Protein degradation by ATP dependent proteases is necessary for short-lived regulatory proteins and abnormal polypeptide. In eukaryotes the most important of these proteases is the proteasome. The proteasome itself is a cylindrical structure of around 700 kDa. The proteolytic sites are in the central core with access through channels of order 13 Å in diameter at their narrowest point. Other ATP dependent proteases, namely ClpXP and ClpAP, complete the same function in organelles and prokaryotes. Substrate proteins are targeted to these proteases by means of N or C terminal targeting sequences. The ability to unfold a protein targeted for degradation is determined by the local structure of the protein next to the targeting signal. Protein tends to be more easily unfolded when the degradation signal leads to the surface of a domain with *α*-helical structure or unstructured loops, while if the signal is found next to buried *β*-strands, unfolding is more difficult to initiate.

Many research group has discussed the controlled unfolding and translocation of proteins through the *α*hemolysin pore using AAA+ unfoldase ClpX [24]. The single *α*-hemolysin pore was inserted into a lipid bi-layer between two compartment (cis and trans). A patchclamp amplifier applied a constant 180 mV potential between two Ag/AgCl electrodes and recorded ionic current through the nanopore as a function of time. Substrate proteins were added to the cis solution and linked ClpX variant was present in the trans solution. ClpX served as a molecular machine that used chemical energy derived from ATP to pull the protein through the pore [25–33]. AAA+ unfoldases denature and translocate polypeptide into associated peptidases. ClpX (an unfoldase) generates mechanical force to unfold its protein substrate from Escherichia coli, alone, and in complex (with combination) with the ClpP peptidase. ClpX hydrolyzes ATP to generate mechanical force and translocate polypeptide chain through its central pore [22, 29, 38, 39]. AAA+ unfoldases target structurally and functionally diverse proteins in all cells. Also, AAA+ target proteins not only in a folded, soluble conformation, but also in hyperstable mis-folded or aggregated states. These molecular machines must use efficient mechanisms to unravel proteins with a wide range of thermodynamic stabilities, topologies, and sequence characteristics. These results shows that ClpX(P) is able to generate and apply mechanical forces sufficient to unfold most target proteins.

In recent past, some theoretical and experimental work has been done where the width of the pore changes during the translocation process. In such translocation process, the dynamical nature of the pore enable the polymer chain to translocate more efficiently as compare to the translocation of polymer in static pore [40]. On the experimental side, it has been shown that the translocation of DNA through a nanochannel can be modulated by dynamically changing the cross section of an elastomeric nanochannel device by applying mechanical stress [41–45]. Time-dependent driving force are also used in the translocation process [46–50]. There are some biological examples of such fluctuating environment in translocation are the nuclear pore complex, which plays an essential role in nucleocytoplasmic transport in eukaryotes [51]. Another example is the exchange of molecules between mitochondria and the rest of cell which is controlled by the twin-pore protein translocate (TIM22 complex) in the inner membrane of the mitocondria [52]. Moreover, using an alternating electric field (time-dependent driving force) in the nanopore has been implied as a source for DNA sequencing [53].

P. Fanzio, et al., [42] has shown that the DNA translocation process through an elastomeric nanochannel device can be altered by dynamically changing its cross section. In such experiment, the deformation in nanochannel is induced by a macroscopic mechanical compression of the polymer device. It has been demonstrated that it is possible to control and reversibly tune the direction of a nano-channel fabricated using elastomeric materials, so to fit the target molecule dimensions. The single DNA molecules are electrically detected at different squeezing conditions, which show the possibility to manipulate the DNA passage inside the nano-channel; that is the translocation time of DNA molecule obtained at high applied displacement is an oder of magnitude higher compared to the one obtained without applying the compression. The opportunity to dynamically control the nano-channel dimension open up new possibilities to understand the interactions between bio-molecules and nano-channels, such as the dependence of the translocation dynamics on the channel size, and the effects of moving walls [40].

Here we discuss the translocation of a protein through a flickering pore which is characterized by measuring the average translocation time as a function of the importing force *F*. The average was performed over independent runs, excluding those in which protein did not cross the membrane channel from the CIS to the TRANS-side within the waiting time *τ*_*w*_ = 10^5^*t*_*u*_. The average translocation time grows with the decrement of external force *F*, indicating that translocation processes is drastically slowed down. Which indicates the presences of a free-energy barrier due to channel-protein interaction, which the molecule has to overcomes to activate its translocation process. The comparison reveals that at the limit of low external force *F*, the translocation of protein through flicking pore takes short time as compared to its static counterpart. The difference between the curves representing the translocation times are remarkable particularly at the extreme limit of low force. We note that a pore with fluctuating radius enhances the translocation process as compared to the static pore that has a radius equal to the mean radius of the fluctuating pore. The two curves overlap at the limit of high force due to fast translocation process, hence the static and fluctuating pore becomes indistinguishable.

The paper is organized as follows: sec. II we derscibe the computational model used to simulate GFP unfolding in a confinement cylindrical geometry. We used coarse grained Go model both the protein and confinement geometry. In sec.III we discuss the results of protein unfold and translocation through a dynamical nanopore. Sec. IV is devoted to the conclusion.

## II. MODEL AND SIMULATION DETAILS

### A. Protein and nanopore model

We used a Gō-like force field acting on the beads in our numerical simulation which preserve the secondary structure content, the beta sheets and alpha helices, of a protein chain [56–61]. In this approach the total potential acting on all the residues of the proteins is [62–64]:

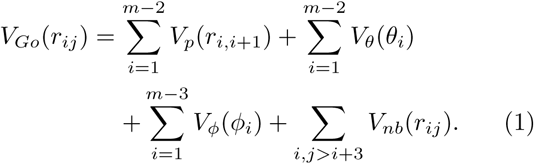

where peptide potential *V*_*p*_(*r*_*i*,*i*+1_), responsible for the covalent bonds between the beads of the polymer chain, has the following expression:

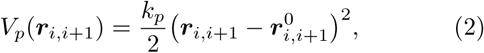

where ***r***_*i*,*i*+1_ = ***r***_*i*_ − ***r***_*i*+1_, 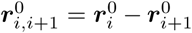 are the position vector between the bonded residues *i* and *i*+1 in the instantaneous and native configuration respectively. The norm of position vector is the bond length. The empirical constant 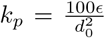 is in term of the equilibrium length parameter *d*_0_, and is in the unit of energy parameter *ϵ*. In our simulation, *d*_0_ = 3.8 Å is the average distance between two adjacent amino acids and *ϵ* sets the energy scale of the model. The angular potential *V*_*θ*_ is used to recover the secondary structure of protein in reference native conformation. Mathematically,

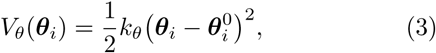

where *k*_*θ*_ = 20*ϵ* rad^−2^ is the elastic constant expressed in term of the energy computational unit *ϵ*, and ***θ***_*i*_, 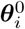 are bond angles formed by three adjacent beads in the simulated (time-dependent) and native conformation, respectively.

The dihedral potential (torsion) *V*_*φ*_(*φ*_*i*_) is define as:

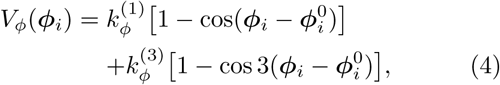

where 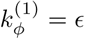 and 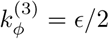 are dihedral constants expressed in term of energy unit *ϵ*.

Non-bonded (nb) interactions between nonconsecutive amino acids are modeled with Lennard-John 12-10 potential.. The expression for Lennard-Jones potential is:

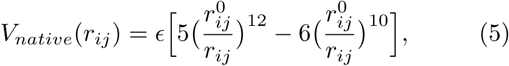

where all the symbols have been already defined. When 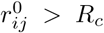, the purely repulsive contribution *V*_*nonnative*_ is assigned to the pair of amino acids considered. This term enhances the cooperatively in the folding process and takes the form of a Lennard-Jones barrier

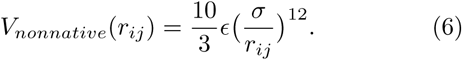

The non-bonded potential *V*_*nb*_ summarized the possible long range interaction just described above and reads as

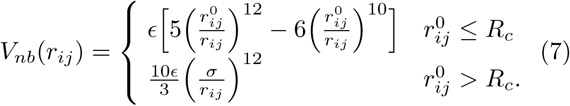

The confinement effect on protein dynamics can be represented by a step-like soft-core repulsive cylindrical potential. The expression of the pore potential is given by:

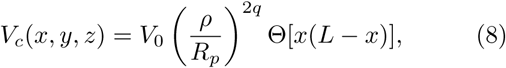

where *V*_0_ = 2ϵ and Θ(*s*) = [1 + tanh(*αs*)]/2 is a smooth step-like function limiting the action of the pore potential in the effective region [0,*L*]. *L* and *R*_*p*_ are pore length and radius respectively. Also, 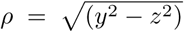 is the radial coordinate. The parameter *q* tunes the potential (soft-wall) stiffness, and *α* modulates the soft step-like profile in the *x*-direction; the larger the *α*, the steeper the step. In our simulation, we consider *q* = 1 and *α* = 2 Å^2^. The driving force *F*_*x*_ acts only in the region in front of the pore mouth *x* ∈ [−2, 0], and inside the channel [0, *L*]. Pore length *L* = 100 Å and radius *R*_*p*_ = 10 Å are taken from *α*HL structure data.

### B. Equation of Motion: Langevin dynamics

The equation of motion which governs the motion of material point is the well-know Langevin equation. The Langevin equation is given by

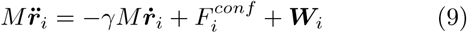

where 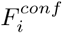 is the sum of all the internal and external forces acting on residue *i*. Here *γ* is the friction coefficient used to keep the temperature constant (also referred as Langevin thermostat). The random force ***W***_*i*_ accounts for thermal uctuation, being a delta-correlated stationary and standard Gaussian process (white noise) with 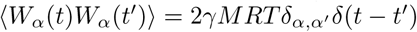

## III. FLICKERING NANOPORE

In order to produce fluctuations in the pore, the confinement effect on a protein driven inside the narrow space such as a nanopore can be modified by introducing a periodic oscillation in the repulsive cylindrical potential. This can be done by making the radius of cylinder time dependent. Hence, the pore potential through which proteins are imported becomes

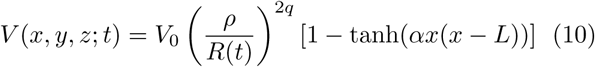

where *R*(*t*) = *R*_*P*_ (1 − *R*_*A*_ sin(*ωt* + *θ*)) is the time dependent radius of the repulsive cylindrical potential. The *R*(*t*) allows the radius of the pore to oscillate harmonically, with frequency *ω* = 2*πT* and amplitude *R*_*A*_, about an average width *R*_*p*_. This defines the *R*_*min*_ = *R*_*p*_ − *R*_*A*_ and *R*_*max*_ = *R*_*p*_ + *R*_*A*_, the minimal and maximal radii of the pore, respectively. The *θ* represents the initial phase.

## IV. RESULTS

## V. GREEN FLUORESCENT PROTEIN (GFP)

Green fluorescent protein (GFP) [65] from the jellyfish Aequorea victoria is one of the most important proteins currently used in biological and medical research having been extensively engineered to be used as a marker of gene expression and protein localization, as an indicator of protein-protein interactions and as a bio-sensor [71]. The GFP having 238 residue undergoing an auto-catalytic post-trans-locational crystalization and oxidation of the polypeptide chain around residues Ser65, Tyr66 and Gly67, to form an extended conjugated *π*system, the chromophore, which emits green fluorescence [72].

The remarkable cylindrical fold of the GFP seems ideally suited for its function. The strands of the *β*-sheet are tightly fitted to each other like stave in a barrel and form a regular pattern of hydrogen bonds, as shown in Fig. (1). Together with the short *α*-helices and loops on the ends, the can-structure forms a single compact domain and does not have obvious clefts for easy access of diffusible ligands to the fluorophore. The florescence properties of GFP depend on its folded structure. For example, denatured GFP display very low tolerance because of solvent quenching. In native GFP, by contrast, an 11-stranded *β* barrel shields the enclosed chromophore (residue 65:67), which contains a phenolic side chain that equilibrates slowly between protonated and unprotonated states [66–68].

**FIG. 1.**
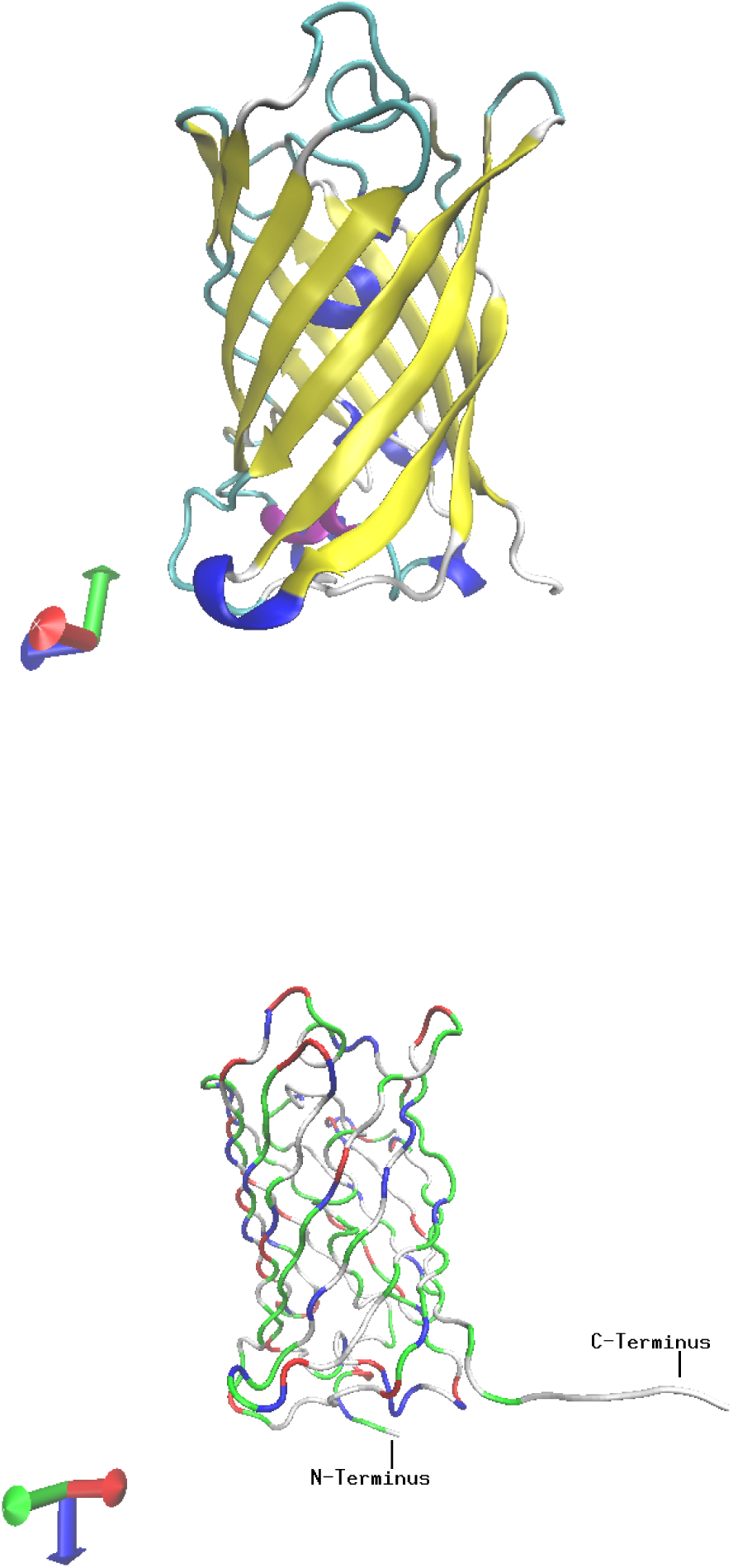
The overall shape of the GFP and its association into dimers. Eleven strands of *β*-sheet (yellow) form the walls of a cylinder. Short segments of *α*-helices (blue and purple) cap the top and bottom of the *β*-can and also provide a scaffold for the fluorophore, which is near geometric center of the can. The *β*-sheet outside and the helix inside, represent a new class of proteins. GFP with colors that vary according to the residues type. The N and C-termini are marked. To facilitate the trapping inside the confinement geometry i.e cylinder, a linker without structure are added to the C terminus. Figure produced by VMD-software.

**FIG. 2.**
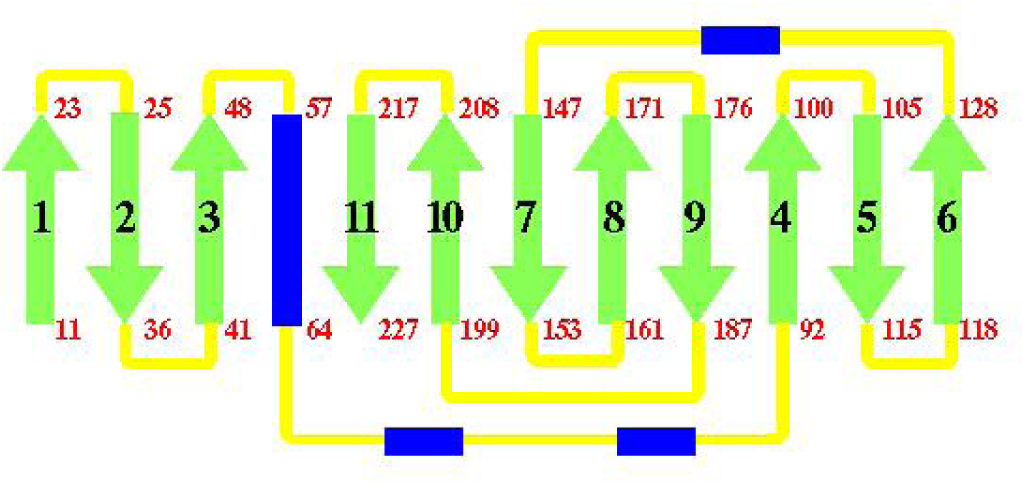
A topology diagram of the folding pattern in GFP. The *β*-sheet strands are shown in green color, *α*-helices in blue, and connecting loops in yellow. The position in the sequence that begin and end each major secondary structure element are also shown. The anti-parallel strands (except for the interactions between strands 1 and 6) make a tightly formed barrel.

The unprotonated chromophore absorbs 467-nm light and emits 511-nm fluorescence. The protonated chromophore absorbs 400-nm light but also emits a 511-nm photon, because absorption transiently leads to deprotonation through an excited-state proton transfer reaction [69, 70].

## VI. UNFOLD OF PROTEINS THROUGH *α*-HEMOLYSIN PORE

Figure (4) describes the plot of typical unfolding trajectory as a function of time of GFP model when the force is applied to the C-terminal. Based on the topology map of GFP, an unfolding intermediate with the N-terminal residues still folded would require part of the beta-strand to be unstructured. As shown in Fig. (3), the folded intermediate stage enter as a loop inside the channel and translocates in double file conformation. Unfolding trajectory of the GFP molecule shows mainly one or two rips, each one followed by translocation of unfolded polypeptide chain [37].

**FIG. 3.**
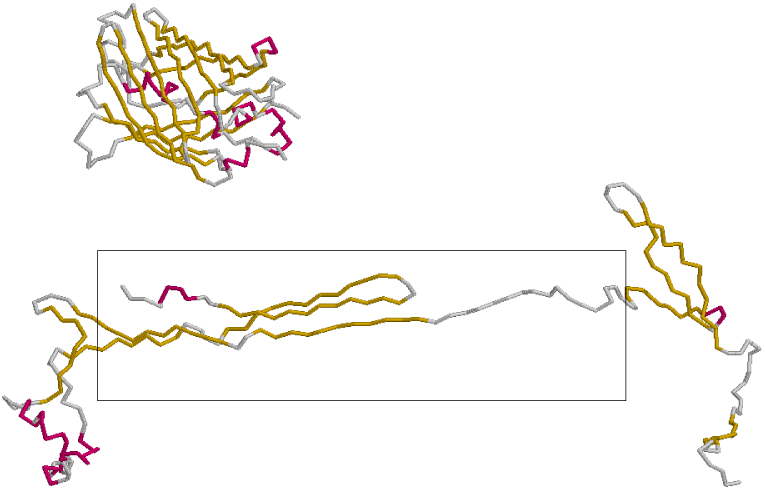
A schematic diagram of GFP unfolding and translocate through the *α*-hemolysin. The upper panel shows the GFP molecule before unfolding. The bottom panel shows the unfolding intermediate and its translocation through pore.

**FIG. 4.**
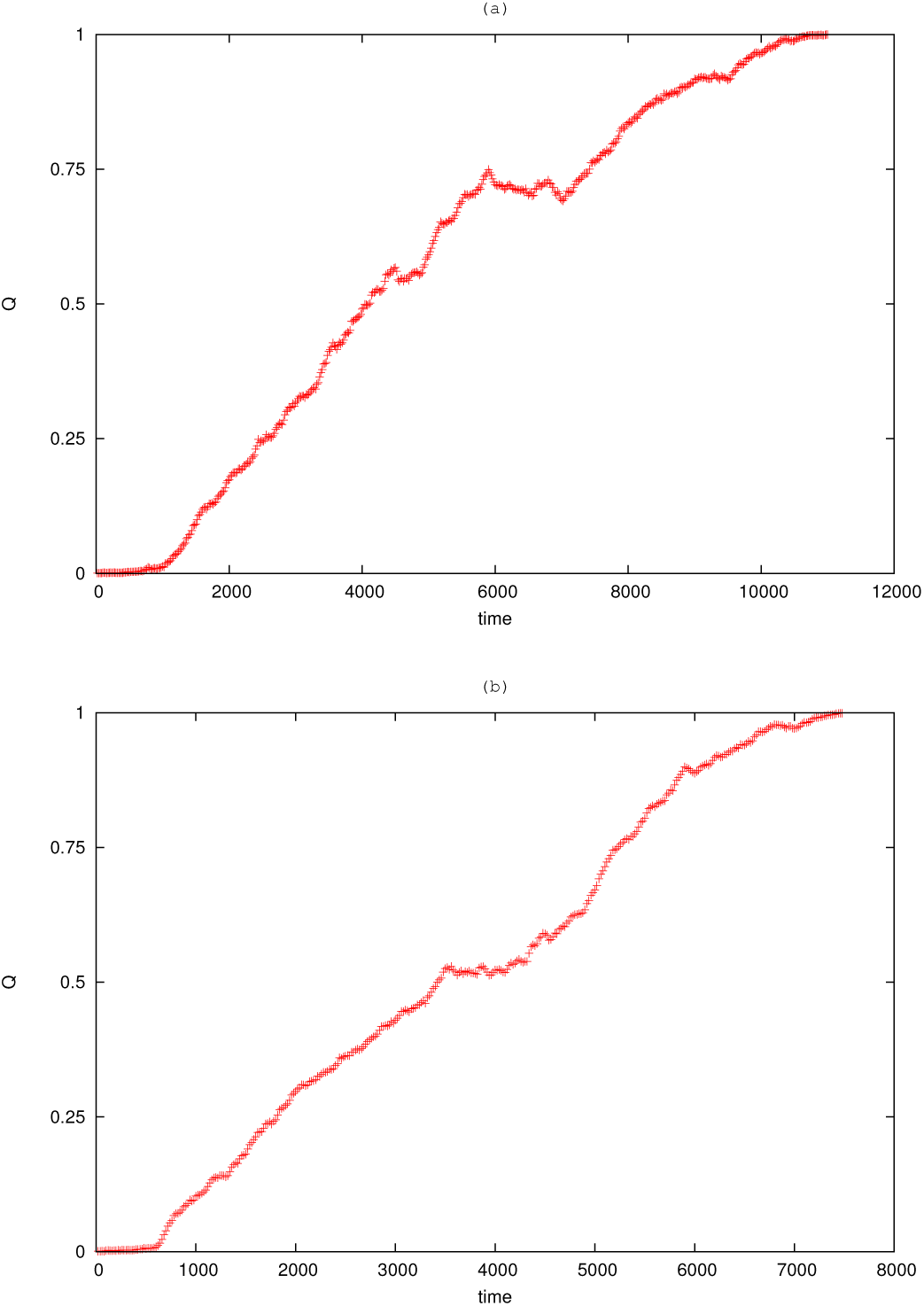
Trajectory of GFP in collective variable coordinate, define in [76, 77], as a function of time during unfolding and trans-location through a static *α*-hemolysin pore effected using a constant external force of (a) *F* = 1.5, and (b) *F* = 2.0. at the C-terminus. The rip in the trajectory corresponding to a GFP unfolding is preceded by a pause or stall effect. After GFP is unfolded, the pore trans-locates the unfolded polypeptide chain with occasional pauses.

## VII. TRANSLOCATION OF PROTEINS THROUGH FLICKERING PORE

We consider the translocation of GFP molecule through a flickering pore. We consider the case where the radius of the pore is time dependent *R*(*t*) = *R*_*p*_(1 − *R*_*A*_ sin(*ωt*)), where *ω* is the frequency of pore oscillation, *R*_*A*_ is the amplitude, and *R*_*p*_ is the average radius of the channel. This modification in pore potential allows the channel to oscillate harmonically with certain frequency. We used initially a linker in the GFP to help the first residue fixed at the entrance of the pore while the other residues are allowed to fluctuate.

When the simulation starts, the first residue of the GFP (first monomer in polymer case) is released under the action of driving external force *F* and the time that elapses between the entrance of this residue into the pore and the exit of the last residue is measured. This gives us the first passage time of the GFP through the pore.

Figure (5) shows the distribution of translocation time. In the limit of high and low flickering rate of pore oscillation, the time distribution is very similar to the zero amplitude case. The green dotted line reveals the distribution when the frequency is *ω* = 0: translocation through static pore. In general, the distribution in static case is Gaussian with an exponentially decaying tail. At the intermediate regime of pore oscillation, the distribution consists of a series of peaks separated by distinct minima. These minima correspond to the points in the pore oscillation cycle when the width of the pore is at its minimum, *R*_*min*_.

**FIG. 5.**
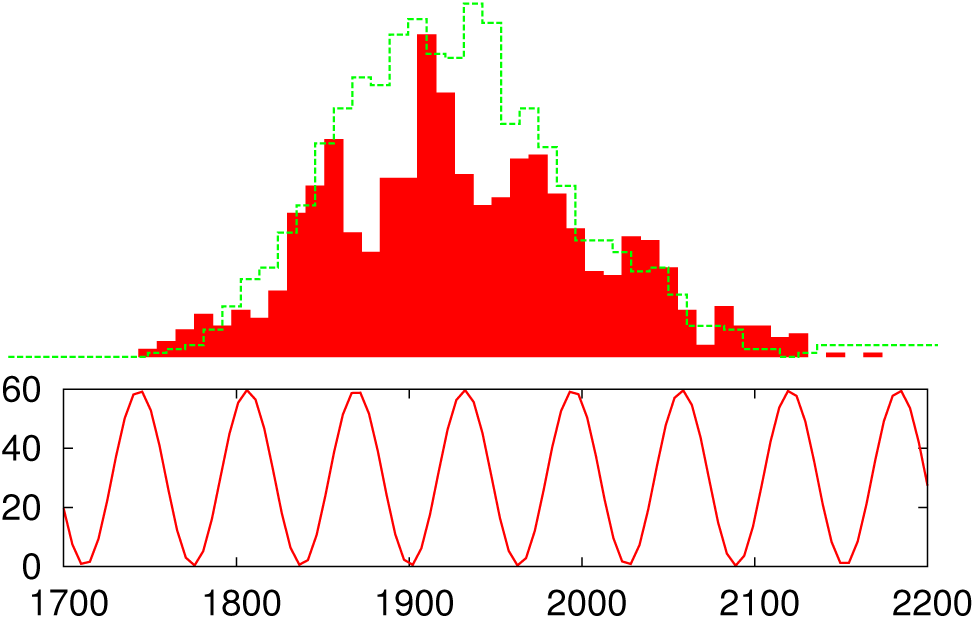
Distribution of first passage time when trans-locating GFP through a flickering pore at frequency *ω* = 0.1 and external force *F* = 5.0. The translocation time distribution are compared with the distribution for the static pore with average radius (green dotted line). The matching of the pore frequency oscillation are also shown to match with the peaks within the time distribution..

We study the effect of changing the amplitude *R*_*A*_ of the oscillating pore on the translocation time. The result for the translocation time as a function of frequency *ω* are shown in Fig. (7). With this particular choice of amplitude *R*_*A*_ = 0.3 Å, the radius of the cylindrical pore oscillate harmonically with minimal radius *R*_*min*_ = 7 Å and maximal radius *R*_*max*_ = 13 Å about an average radius *R*_*p*_ = 10 Å. We find a one global maxima followed by a series of local maxima and minima as illustrated in Fig. (7). The gain in translocation time defined as *ζ* ≡ *τ*/*τ*_0_, where *τ*_0_ represents the time in the static case, is plotted as a function of dimensionless frequency *ω*/*ω*_0_. As compared to the static pore, the translocation rate increases for the oscillating pore. As a function of frequency *ω*, we observed three distinct regions for the translocation rate. In the limit of high frequency, the translocation gain becomes insensitive to the frequency. For small frequencies the average translocation time increases steadily and develops a plateau, as shown in the inset of Fig. (7). During the first half period of pore oscillation, the radius of the pore is greater than the width of the static pore, *R*_*p*_. The translocation time increases for low frequency because the protein only feels this half of the cycle before the completion of the translocation process. During the next half cycle where the pore radius is smaller than the static pore radius, the protein experiences this cycle and as result the gain decreases until reaches a minimum. When the frequency tends to zero, the pore is static and the protein translocation time reduces to the translocation time for the static pore and the gain approaches unity. At the intermediate regime of pore oscillation, the oscillation gets rapid. In this particular phase of pore oscillation the protein either spends substantial amount of time in trapped state or translocate very fast, as shown in the scatter data of translocation time at *ω* ~ 0.3, Fig. (6).

**FIG. 6.**
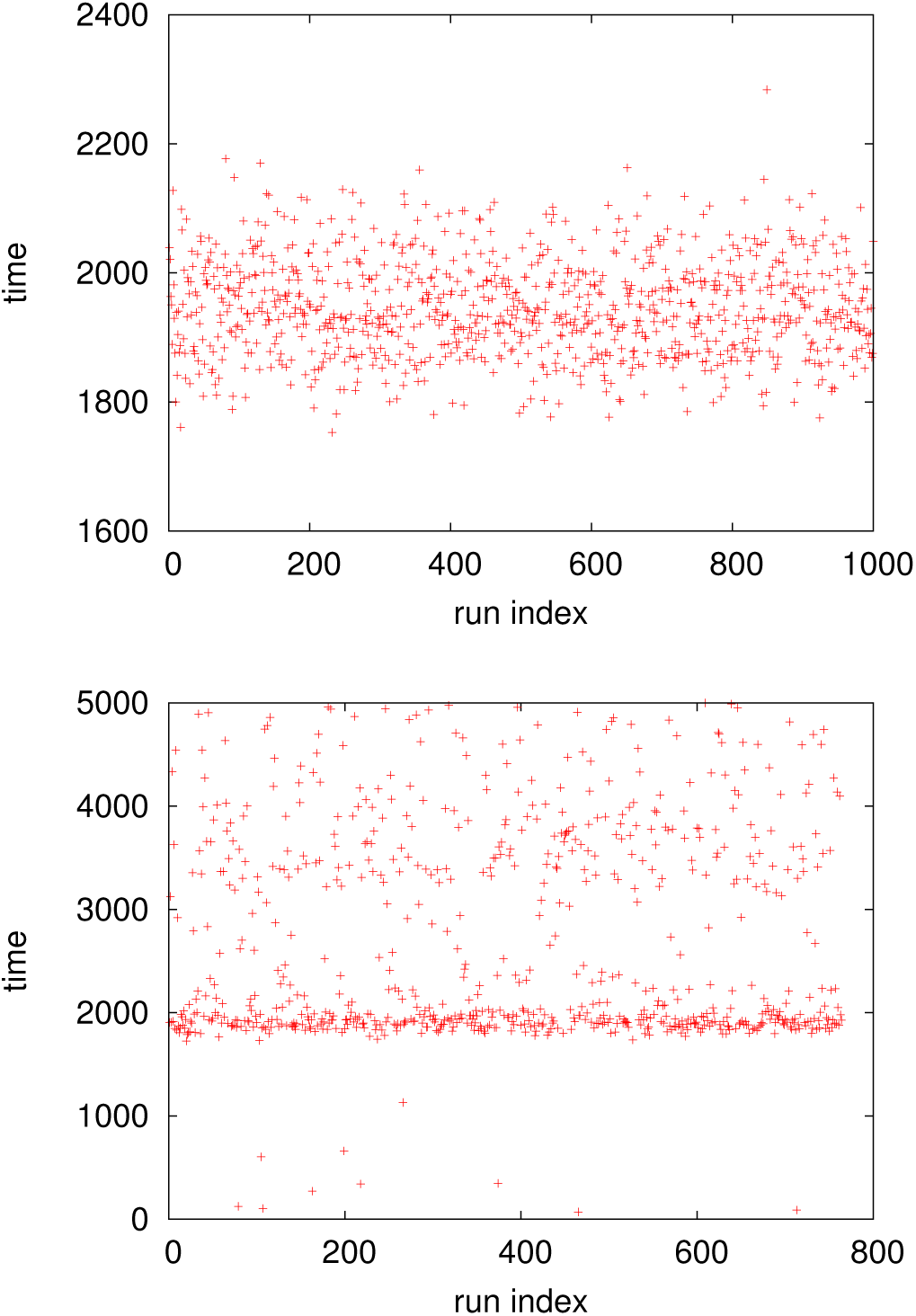
Top Panel: Scatter data of translocation time of GFP at *ω* = 0.1. Lower Panel: Scatter data of translocation time at *ω* = 0.3. The data at *ω* = 0.3 are more scattered and noisy subjected to the fast oscillation of the channel.

**FIG. 7.**
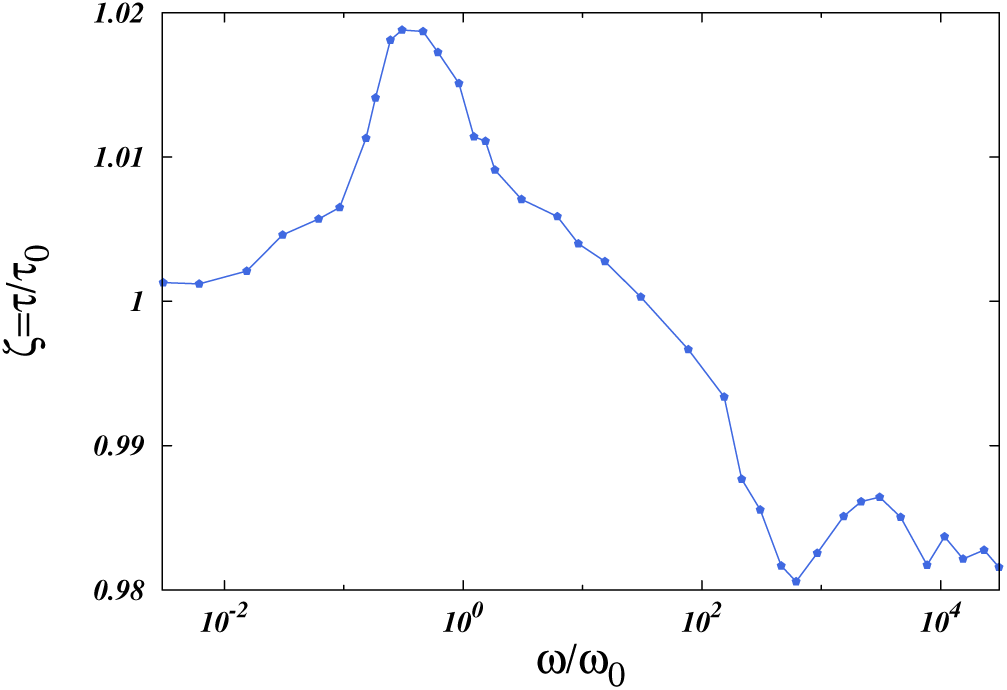
Average gain in translocation rate as a function of dimensionless frequency *ω*/*ω*_0_[*ω*_0_ = 2*π*/*τ*_0_] for aptitude *R*_*A*_ = 0.3 using GFP molecule.

The gain increases initially due to translocation of protein through a narrow opening of the pore. However, during the next half cycle of pore oscillation where the pore radius is larger than the static pore radius *R*_*p*_, the protein only experiences this half of the cycle before the completion of translocation time. As a result the gain in translocation rate decreases due to large pore opening. When the frequency is further increased, the gain becomes relatively insensitive to the frequency and develops a plateau. As the frequency tends to zero *ω* → 0, the gain approaches unity because the pore is basically static and the protein translocation time reduces to the translocation time for static pore. The amplitude plays an important role in the translocation process due the large size of the pore i.e *L* = 100 Å. With large length of pore, the fluctuating phenomena either trap the protein for substantial amount of time or translocation it very fast as already demonstrated in Fig. (7) along with the scatter Fig. (6).

## VIII. *β*-BARREL AND *α*-HELICAL MEMBRANE PROTEINS

We consider the unfolding and translocation processes of the following two different proteins through static pore.

- Green fluorescent protein (*β*-barrel or *β*-can protein) consists of 11 strands of *β*-sheet with an *α*-helix inside and short helical segments on the ends of its cylindrical shape.
- Maltose Binding protein (*α*-helical protein) is folded from a single polypeptide chain of 370 residues. The maltose-binding protein consists of 40% *α* helix and 20% *β* sheet.

Two different types of protein, one with majority of *β*-sandwich proteins (GFP) [37] and another with majority of *α*-helix proteins (MBP) [73], show significant different behavior when unfold and translocate through a channel by external force. We find a conformation of *β*-hairpin when translocating GFP through a static pore. As shown in the topology structure of GFP in Fig. (2), the anti-parallel *β*-strands make a tightly formed barrel. On the other when force applied parallel to the hydrogen bonds leads each bond in turn, resulting a peeling rupture of each bond at relatively low force.

Unfolding and translocation of MBP mechanically is in contrast with the GFP molecule. The long polypeptide chain of MBP unfold and translocation in single file conformation. The MBP unravel begins at the C-terminus where the three *α*-helices are mechanically unravel at very low external forces. On the other side, the N-terminal domain which consists of a five *β*-strands unravel at the expenses of high force (the configuration of *β*-sheets and *α*-helices are illustrated in Fig. (8)) [79].

**FIG. 8.**
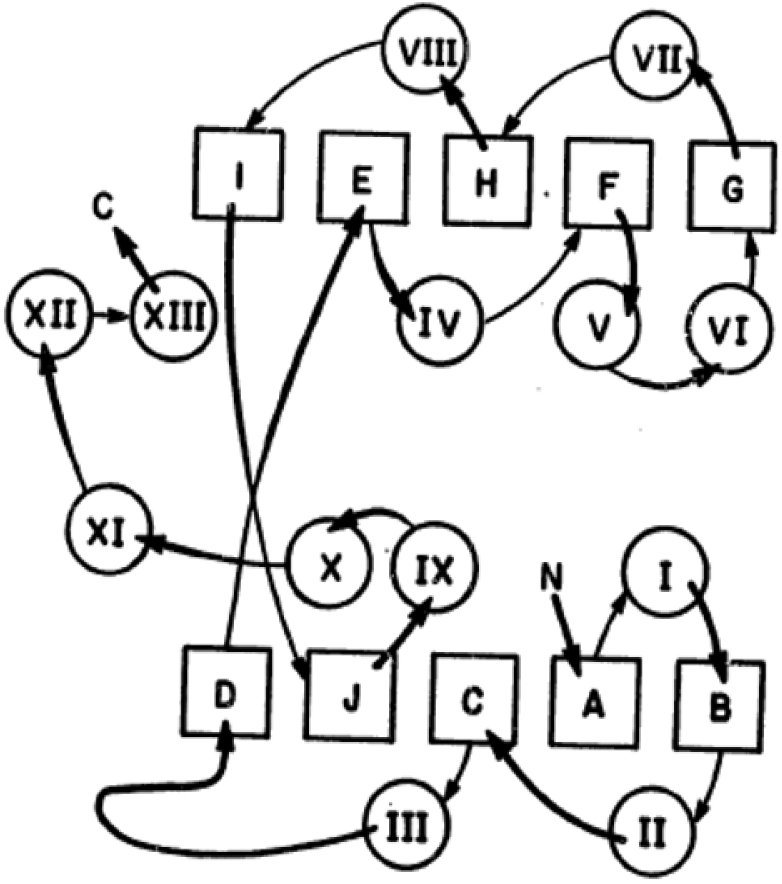
A topological packing diagram for the maltosebinding protein. The squares represent strands in *β*-sheets and are designated by letters. The circles labeled with Roman numbers represent *α*-helices. The N- and C-terminal are also labeled in the figure.

Figure (9) shows the trajectory of center of mass coordinate (reaction coordinate *X*_*CM*_) as a function of simulation time for the GFP molecule (red empty dots), and for the MBP molecule (green filled dots) when translocating through a static pore. The rip in the trajectory corresponding to a GFP unfolding is preceded by a long pause or stall effect as compared to the short rip in the MBP trajectory. MBP takes longer time in unfolding but once it overcomes the entropic barrier it translocates with a small rip [78].

**FIG. 9.**
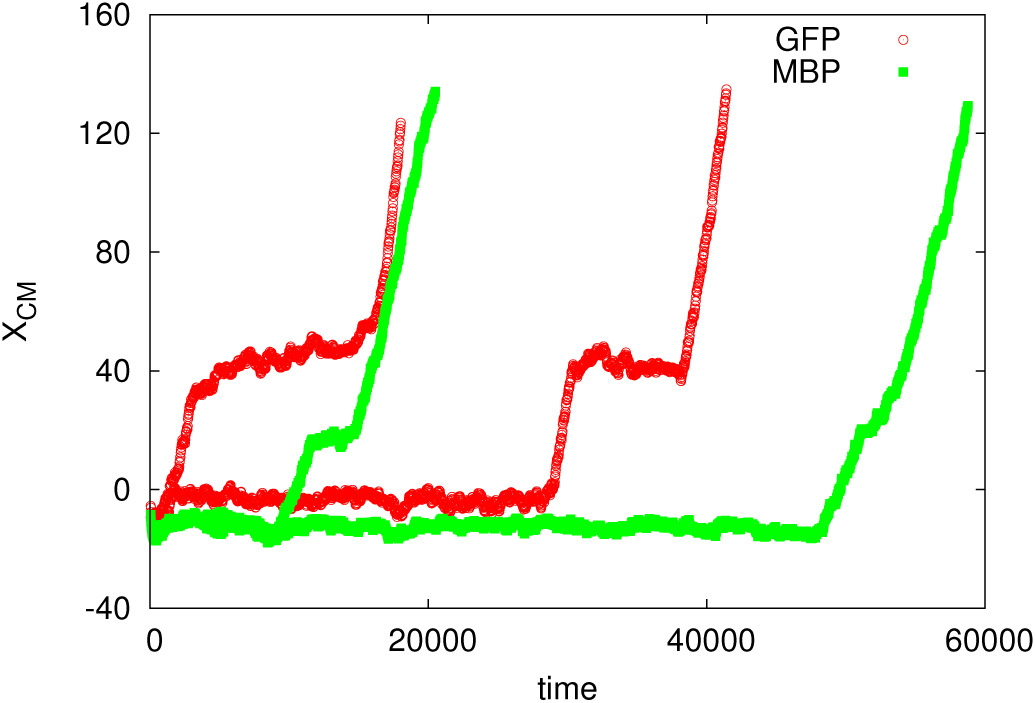
Translocation reaction coordinate as a function of time. The red empty dots represent the trajectory of GFP molecule when translocating through a static pore. The green filled dots describe the trajectory of center of mass coordinate using MBP molecule using a constant force *F* = 2.0 at the C-terminus.

## IX. REMARKS ON MBP TRANSLOCATION THROUGH DYNAMIC PORE

In this section, we present a computational study of MBP translocation across a flickering nano-pore to determine how the dynamic pore influence the *β*-sheets and *α*-helices of MBP during its transport. The translocation of MBP confirmation required relatively low force when chemically unfolded, and once one protein terminus (either C or N-terminus) enters the pore, the transport proceeds uniformly without further resistance from the free-energy barrier. However, native-like MBP conformation exhibit a much richer translocation phenomenology; large force are required to trigger the translocation process. The translocation of native-like MBP conformation proceeds through bottlenecks and jerky movements due to the structural rearrangements of the folded part of the protein that has not yet engaged the pore (as illustrated in the trajectory Fig. (9, green filled dots)). The transport of MBP across the pore temporarily stalled at the well-defined stage of translocation of process. The stall effect is more dominant in case of GFP which consists of majority *β*-sheets. Hence it is very interested to see how the flickering mechanism of pore affect the translocation process of MBP.

Figure (10) shows the distribution of MBP translocation time at difference frequency of pore oscillation. The scatter data of translocation time (red dots in Fig. (10)) is more noisy at low frequency i.e *ω* = 0.5, 0.3. We observed more organized data of translocation time at relatively high frequency. It is very informative to inspect the distribution of translocation time at different frequency. As shown in Fig. (10), we find that the distribution consists two peaks at *ω* = 0.3 separated by distinct minima. These minima correspond to pore oscillation when the radius of the pore is at its minimum. When the frequency of pore oscillation is further increased, it causes an increase in the numbers of peaks in the translocation time distribution. Even though the locking of peaks in the distribution of translocation time with frequency of the pore oscillations is only approximated but this partial dephasing is probably the signature of the role of a tertiary structure of proteins with respect to linear polymers [40].

**FIG. 10.**
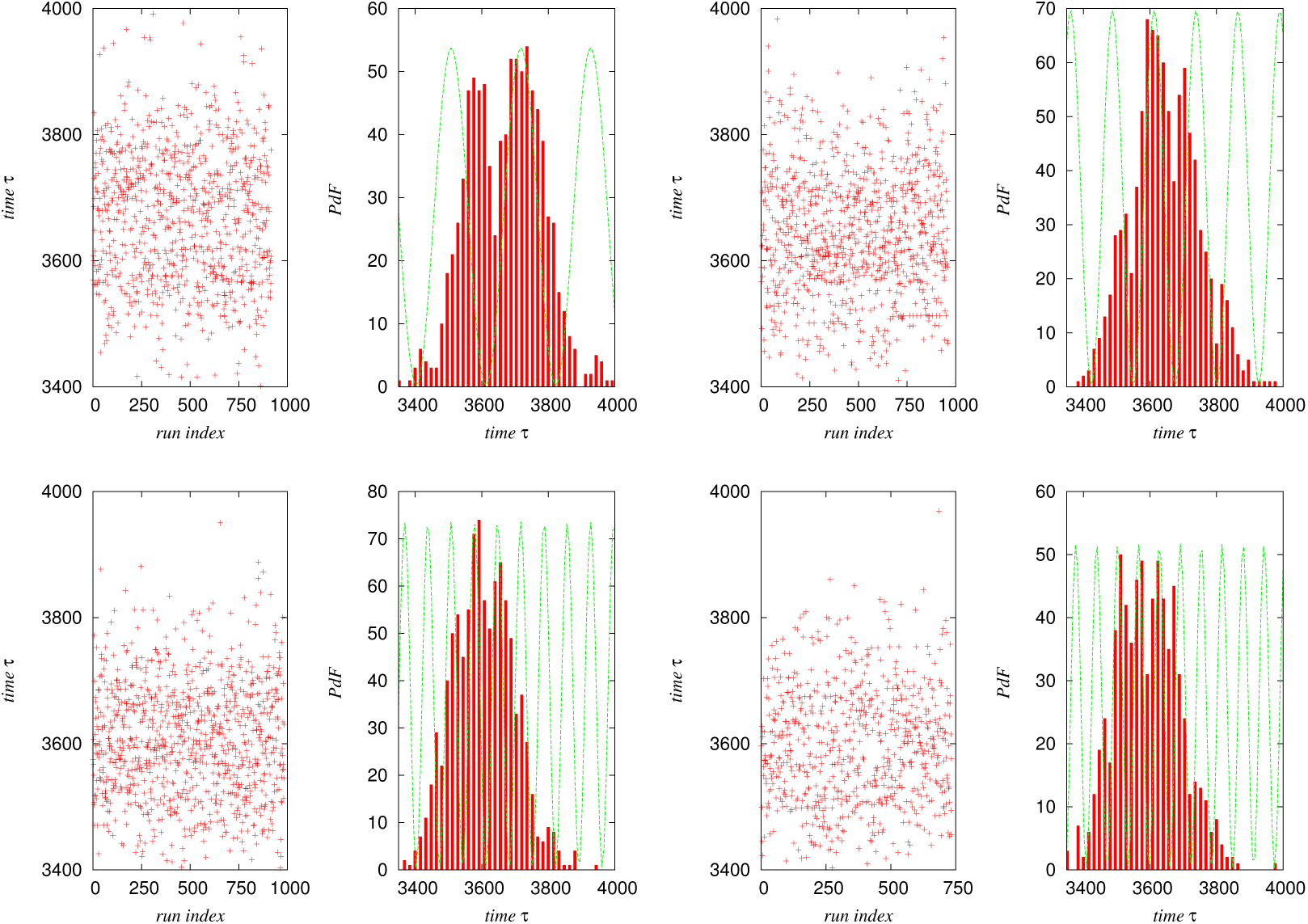
The distribution of MBP translocation time when transport across a flickering pore for *F* = 5.0, *R*_*A*_ = 0.5 Å, *R*_*min*_ = 5.0 Å, and *R*_*max*_ = 10 Å at frequency *ω* = 0.3 (upper left panel), *ω* = 0.5 (upper right panel), *ω* = 0.9 (lower left panel), and *ω* = 0.1 (lower right panel). The red dots represents the scatter data of translocation time and the green dotted line shows the matching of the pore oscillation cycle with the peaks of translocation time distribution.

## X. CONCLUSION

We have simulated Green Fluorescent Protein (GFP) translocation across the flickering *α*HL nano-pore via a coarse grained computational model for both the GFP and the pore. For the sake of comparison, we also run simulations for a polymer equivalent to the GFP molecule in size. This is simply done by setting parameter *R*_*c*_ = 3 Å which determines the number of native attractive interactions and eliminates the interactions which make the protein globular. We simulate the translocation process by allowing the protein cross a free-energy barrier from a metastable state, in the presence of thermal fluctuations. The fluctuation in the channel, which we model by making the radius of pore time dependent, can be originated by the random and non-random (systematic) cellular environment, drive out the protein out of equilibrium during the transport dynamics. The average translocation time as a function of frequency of the pore oscillation reveals a nonmonotonic behavior, characterized by the presence of a global maxima at a specific frequency. Furthermore, we also study the translocation of maltose binding protein (MBP) through flickering pore to show its important topological different from GFP molecule. Translocation of protein across fluctuating environment can play a fundamental role in bio-molecule transport process. The flickering effects of the pore can enhance or speed down the protein translocation depending on the frequency *ω* of the pore oscillation, and on the initial phase *θ*. Hence, the oscillating nature of pore can play the role of tunning mechanism to select a particular translocation time of the protein. Our results can be of fundamental importance for those experiments on DNARNA sorting and sequencing and drug delivery mechanism for anti-cancer therapy.

The above studies of GFP and MBP proteins shows that the transport of long structured bio-molecules (proteins and DNA-RNA nucleic acids) across nano-scale channels is a complex process which is achieved via several structured rearrangements such as the unfolding of structured protein at the cis-side of the pore, the effects of flickering pore on translocation dynamics and the refolding of protein at the trans-side of the pore. In this paper we give a description of the complex phenomenology observed in experiments on biomolecules translocation requires an integrated approach able to combine the molecular dynamics simulations, statistical analysis of dynamics, and techniques adopted from stochastic processes theory.

## ACKNOWLEDGMENTS

The author would like to thank U. M. B. Marconi, A. Vulpian and F. Cecconi for helpful discussion, and Institute of Complex Systems, National Research council of Rome for providing access to computational resources.

